# A high affinity human monoclonal antibody against Pfs230 binds multiple parasite stages and blocks oocyst formation in mosquitoes

**DOI:** 10.1101/2020.09.25.313478

**Authors:** Camila H. Coelho, Wai Kwan Tang, Martin Burkhardt, Jacob D. Galson, Olga Muratova, Nichole D. Salinas, Thiago Luiz Alves e Silva, Karine Reiter, Nicholas J. MacDonald, Vu Nguyen, Raul Herrera, Richard Shimp, David L. Narum, Miranda Byrne-Steele, Wenjing Pan, Xiaohong Hou, Brittany Brown, Mary Eisenhower, Jian Han, Bethany J. Jenkins, Justin Yai Alamou Doritchamou, Margery G. Smelkinson, Joel Vega-Rodriguez, Johannes Trück, Justin J. Taylor, Issaka Sagara, Jonathan P. Renn, Niraj H. Tolia, Patrick E. Duffy

## Abstract

Malaria elimination requires tools that interrupt parasite transmission. Here, we characterized B cell receptor responses among Malian adults vaccinated against the first domain of the cysteine-rich 230kDa gamete surface protein Pfs230^1–3^ to neutralize sexual stage *P. falciparum* parasites and halt their further spread. We generated nine Pfs230 human monoclonal antibodies (mAbs). One mAb potently blocked transmission to mosquitoes in a complement-dependent manner and reacted strongly to gamete surface while eight mAbs showed only low or no blocking activity. This study provides a rational basis to improve malaria vaccines and develop therapeutic antibodies for malaria elimination.

## MAIN TEXT

Malaria eradication is a global priority and will require innovative strategies that, in addition to preventing or controlling human infection, might block parasite transmission through mosquitoes. Sequences of matched heavy and light chain variable regions from single human B cells have been used to identify antibodies generated in response to infection or vaccination and inform vaccinology^4–7^. In this study, we apply this approach to examine human antibodies elicited in response to a transmission blocking vaccine (TBV), that used a Pfs230 fragment as antigen. Pfs230 is present on the surface of *P. falciparum* gametocytes and gametes and mediates binding of exflagellating microgametes to red blood cells, thus parasites lacking this protein cannot bind to red blood cells or further develop into oocysts.^1^ We collected Pfs230 domain 1 (D1)-specific single memory B cells **(Extended Data Fig. 1, Extended Data Fig. 2a)** from eight Malian adults immunized with four doses of Pfs230D1-EPA/Alhydrogel® vaccine (Clinicaltrials.gov NCT02334462) to identify functional monoclonal antibodies elicited in response to a TBV. This vaccine aims to neutralize sexual stage *P. falciparum* parasites by targeting Pfs230, a 230kDa gamete surface protein comprised of fourteen 6-cysteine (6-Cys) domains^1–3^. All samples were chosen from subjects presenting high serum Transmission-Reducing Activity (TRA), measured by the capacity of serum antibodies from immunized subjects to reduce the number of oocysts that develop in mosquitoes fed on in vitro cultured *P. falciparum* gametocytes **(Extended Data Table 1)**.

We obtained 272 VH and 351 VL sequences of B cell receptor (BCR) from Pfs230D1-specific single memory B cells from the vaccinees via amplification and sequencing of the V(D)J region **(Extended Data Fig. 3).** When analysing V gene usage of the BCR sequences, 87.5% of the subjects presented Pfs230D1-specific memory B cells using kappa chains derived from IGKV4-1 **(Extended Data Fig. 2e).** This light chain gene has also been identified in sequences of functional human mAbs obtained in response to other *Plasmodium* antigens^4–6,8^. For the heavy chain, IGHV1-69 was the most commonly expressed gene and detected in 100% (8/8) of vaccinees **(Extended Data Fig. 2f)**. IGHV1-69 is one of the most polymorphic loci of the IGHV gene cluster^9^ and is frequently found in broadly neutralizing antibodies generated in response to influenza haemagluttinin^10,11^.

Nine pairs of BCR sequences were chosen for expression of fully human Pfs230D1 IgG1 antibodies by assessing whether the CDR3 sequences were shared between sorted B cells. This approach identifies identical sequences in multiple B cells from the same subject, indicating that they have been activated in response to vaccination. These nine pairs **(Fig. 1a)** represented distinct heavy and light chain germline genes with an overabundance of IGHV1-18 (N=6), IGHV1-69 (N=3), and IGKV4-1 (N=7). The resulting recombinant antibodies bound to Pfs230D1 antigen **(Figure 1d,e, Extended Data Figure 4).** Competitive epitope binning of the nine mAbs suggested they bind three non-overlapping epitopes in Pfs230D1 **(Fig. 1b)**. LMIV230-01 forms a distinct group (Group 1) and has potent neutralizing activity **(Fig. 1b, c)**. The remaining mAbs do not compete with LMIV230-01 and may form two additional epitope groups. Group 2 and 3 mAbs possess low or no neutralizing activity **(Fig. 1c)**. We therefore focused most of our subsequent analyses on LMIV230-01 and to a lesser extent on LMIV230-02, which demonstrated low functional activity.

**Fig. 1.**
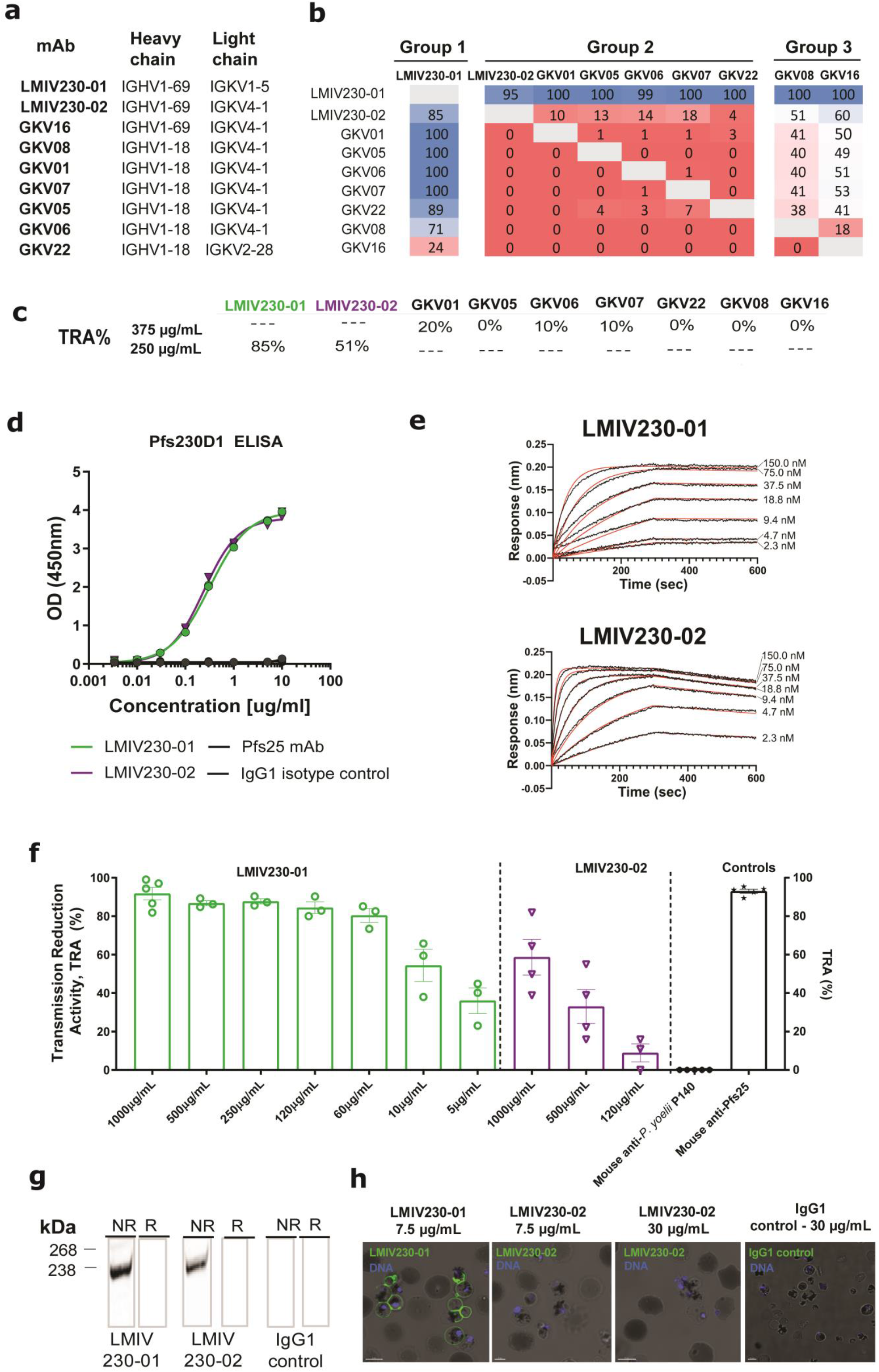
Human recombinant mAbs were generated from Pfs230D1-specific single memory B cells of Malian adults vaccinated with the Pfs230D1-EPA/Alhydrogel® TBV. **a,** VH and VL genes corresponding to each mAb. LMIV230-01 and LMIV230-02 sequences originate from the IGHV1-69 heavy chain gene but utilize different kappa chain genes. Complete V gene usage determined in Pfs230-specific memory B cells is described in **Extended Data Figures 2e,f**. **b,** Epitope binning of human anti-Pfs230D1scFvs. The primary binding scFv is listed on the left and the competing scFv are listed on the top. Reported scores are a percentage of total binding of that antibody in the absence of a competitor scFv. Values greater that 50% display low amounts of competition, while values lower than 50% exhibit greater competition. Any experiment with >100% binding was given a score of 100, while negative values were given a score of 0. Potential epitope bins are grouped and labelled above the table. **c,** Functional activity of each mAb, assessed by Standard Membrane Feeding Assay (SMFA) and measured as the % reduction (versus control mAb) in the number of *P. falciparum* NF54 oocysts in midguts of *Anopheles* mosquitoes (“TRA”). **d,** LMIV230-01 and LMIV230-02 mAbs bound similarly to Pfs230D1 and **e,** show high affinity to recombinant Pfs230D1 **(Extended Data Fig. 4, Extended Data Table 2) f**, LMIV230-01 reduces *P. falciparum* NF54 oocyst numbers by 91.7% at 1000 μg/mL, 86.7% at 500 μg/mL, 84.4% at 250 μg/mL and 80.3% at 60 μg/mL, while LMIV230-02 displays only modest activity with 58.7% reduction at the maximum concentration of 1000 μg/mL, in SMFA. Data from eleven independent SMFA and each concentration was evaluated in at least three biological replicates for each mAb. N ≥ 20 mosquitos per assay. Average oocyst numbers in the control mosquitoes (fed with mouse IgG1 mAb targeting *P. yoelii* P140 protein) for each experiment were: exp. 1 = 29.73; exp.2= 7.18; exp. 3= 57.86; exp. 4= 36.41; exp. 5= 51.71, exp. 6= 4.55; exp. 7= 62.35; exp. 8= 20.50, exp.9= 8.71, exp 10= 18.05, exp. 11= 5.86. Negative oocyst reduction values were set to zero. Human isotype IgG1 and US human serum pool were used as additional negative controls **(Extended Data Fig. 5b)**. Values are shown as mean ± s.e.m. **g,** LMIV230-01 and LMIV230-02 bind to non-reduced (NR) protein extract of *P. falciparum* NF54 gametes purified on Nycodenz after 2 hours in exflagellation medium. **h**, LMIV230-01 strongly binds to gametes at 7.5μg/mL while LMIV230-02 does not bind at 7.5μg/mL. or 30μg/mL. Both mAbs were labelled with Alexa Fluor 488. Scale bars: 5μM.

LMIV230-01 and 02 bound to Pfs230D1 recombinant protein **(Fig. 1d)** with strong and similar binding affinities **(Fig. 1e, Extended Data Fig. 4, Extended Data Table 2).** We confirmed the two mAbs bind distinct epitopes using competition ELISA **(Extended data Figure 5d)** consistent with the epitope binning results **(Fig. 1b)**. Despite their shared use of IGHV1-69, LMIV230-01 and LMIV230-02 displayed numerous differences in their heavy chain CDRs, consistent with their recognition of distinct epitopes **(Extended Data Figure 12)**.

Although presenting similar affinity to Pfs230D1, the mAbs differed in their functional activity as measured by SMFA. LMIV230-01 ablated *P. falciparum* oocyst burden in mosquitoes in a dose-dependent manner with 91.7% neutralization (TRA) at 1000 μg/mL **(Fig. 1f).** Importantly, 80.3% neutralization was retained at 60 μg/mL. On the other hand, LMIV230-02 reduced oocyst burden by only 58.7% at the maximum concentration of 1000 μg/mL and activity was lost at 250 μg/mL. As previously reported, TRA values higher than 80% are highly reproducible across independent experiments^12,13^.Combining the two antibodies did not increase their overall activity: TRA values were not statistically different when 500μg of LMIV230-02 was combined with 10μg of LMIV230-01 (TRA= 58.7%) versus 10μg of LMIV230-01 alone (TRA= 52.5%) in mosquito feeding assays **(Extended Data Figure 5e).**

To understand the differences in functional activity of the two mAbs, we assessed binding to the native protein. Both mAbs reacted to the protein extract of parasites and were sensitive to reduction of the two disulfide-bonds, suggesting the presence of conformational epitopes **(Fig. 1g, Extended Data Fig. 5c**). Interestingly, LMIV230-01 strongly labelled the surface of live *P. falciparum* gametes purified 2 hours post-exflagellation, while LMIV230-02 did not **(Fig. 1h)**. This suggests that the LMIV230-02 epitope is not completely accessible on the surface-displayed native protein, possibly due to structural limitations imposed by the multi-domain protein Pfs230, as has been seen for other proteins^14,15^ including another 6-Cys TBV candidate^16^.

LMIV230-01 bound strongly to fixed parasites in numerous developmental stages including gametocytes, exflagellation centers, microgametes, macrogametes and round forms (zygotes) collected 4 hours after mosquito feeding. As expected, the mAb did not bind to the post-fertilization stage ookinetes, obtained 24 hours after the mosquito bloodmeal **(Fig. 2a)**.

**Fig. 2.**
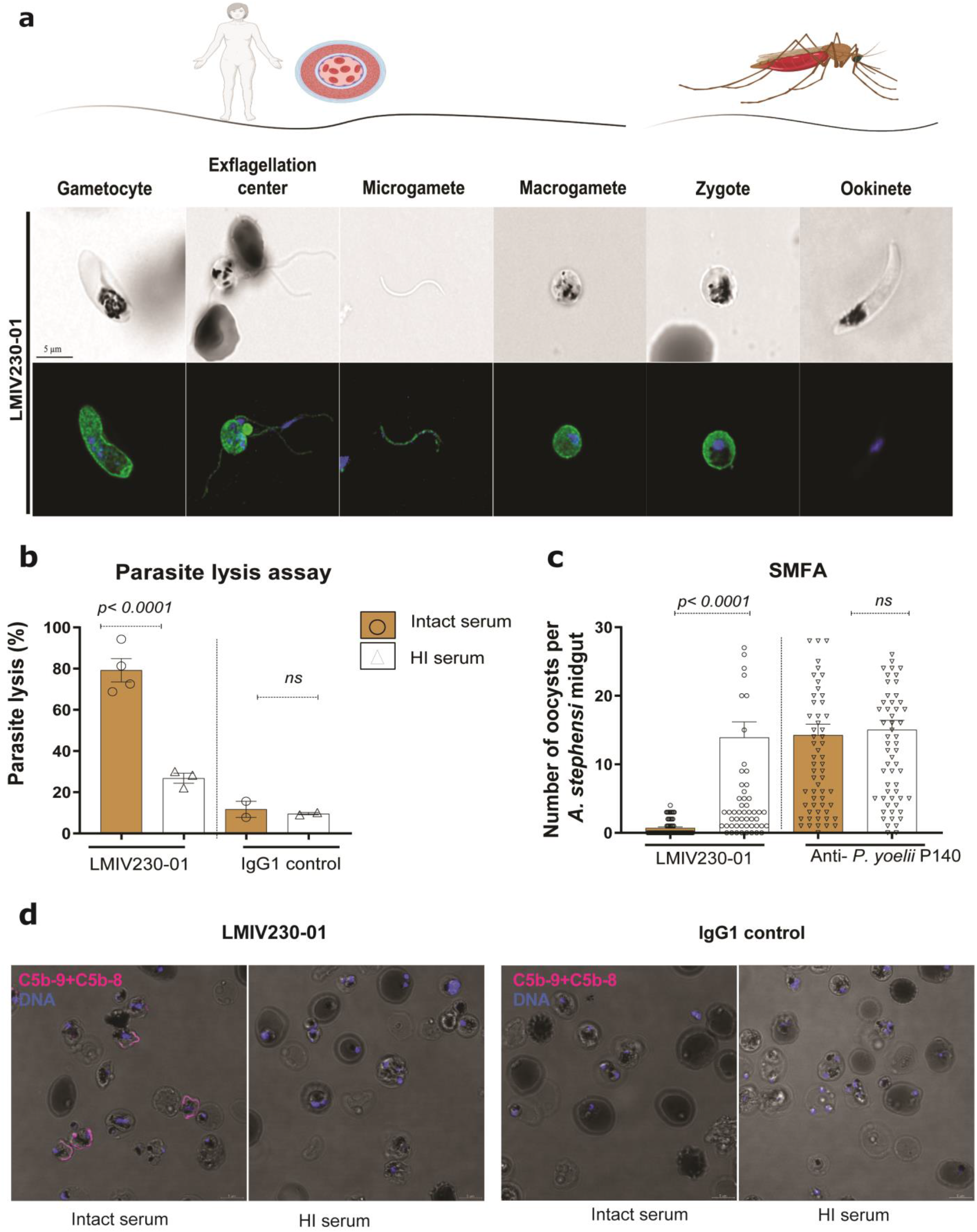
LMIV230-01 binds to multiple parasite stages and its activity is complement-dependent. **a,** LMIV230-01 strongly binds to permeabilized gametocytes, gametes and zygotes and does not bind to ookinetes. Parasites were fixed and permeabilized, and 7.5 μg/mL of antibody was used to stain the different parasite stages. Scale bars: 5μM. **b,** *In vitro* parasite lysis by LMIV230-01 is complement-dependent. Samples were tested in two independent assays, using two different parasite cultures **c,** Functional activity of LMIV230-01 is also complement-dependent *in vivo* (SMFA with mosquitoes). Data from three independent SMFA assays. N ≥ 20 mosquitos per assay. Oocyst averages in the control mosquitoes (fed with IgG1 targeting *P. yoelli* P140) for each of the experiments were: exp. 1= 4.55; exp. 2= 20.50, exp. 3=5.86. Data obtained from mosquitoes fed with LMIV230-01 at 1000 μg/mL with intact sera were also used to generate figure 1f. Values are shown as mean ± s.e.m. One-Way ANOVA and Turkey’s multiple comparisons test were used to compare the different groups **d,** Live imaging of *P. falciparum* NF54 female gametes incubated with LMIV230-01 in the presence of intact serum from a healthy donor revealed surface-deposited MAC (membrane attack complex) using anti-C5b-9+C5b-8 antibody (magenta color). MAC deposition was not detected in the presence of heat-inactivated (HI) serum. Scale bars: 5μM.

Pfs230 antibody activity depends on complement fixation to lyse *P. falciparum*^17^. To test whether the activity of LMIV230-01 was dependent on activation of the complement system, we incubated parasites with LMIV230-01 in the presence of intact or heat-inactivated sera from US donors then assessed lysis of gametes **(Fig. 2b)** as well as transmission of parasites fed to mosquitoes after treatment using the same conditions **(Fig. 2c)**. Functional activity of LMIV230-01 to lyse gametes and block oocyst formation in mosquitoes was substantially reduced in the heat-inactivated sera **(Figs. 2b and 2c),** demonstrating complement-dependency. Activation of complement leads to the formation of the membrane attack complex (MAC), an assembly of the complement molecules C5b, C6, C7, C8, and C9^18,19^ on the parasite surface. Using an antibody that recognizes assembled MAC, we demonstrated complement fixation on the surface of live *P. falciparum* gametes that were incubated with LMIV230-01 in the presence of intact but not heat-inactivated serum **(Fig. 2d and Extended Data Fig. 7).**

To assess whether LMIV230-01 would also bind to other *P. falciparum* strains, we prepared gametocytes from a culture-adapted Malian isolate ^20^ and from St. Lucia strain (originally from El Salvador) ^21^. LMIV230-01 labelled in vitro-induced gametes from both strains **(Fig. 3a and b)**. Induction of gamete stage from the newly characterized Malian isolate was confirmed using a murine anti-Pfs48/45 mAb **(Fig. 3c)**. LMIV230-01 fixed complement on the gamete surface of both strains, confirming that the antibody is functional against heterologous parasites **(Figs. 3d and e).**

**Figure 3.**
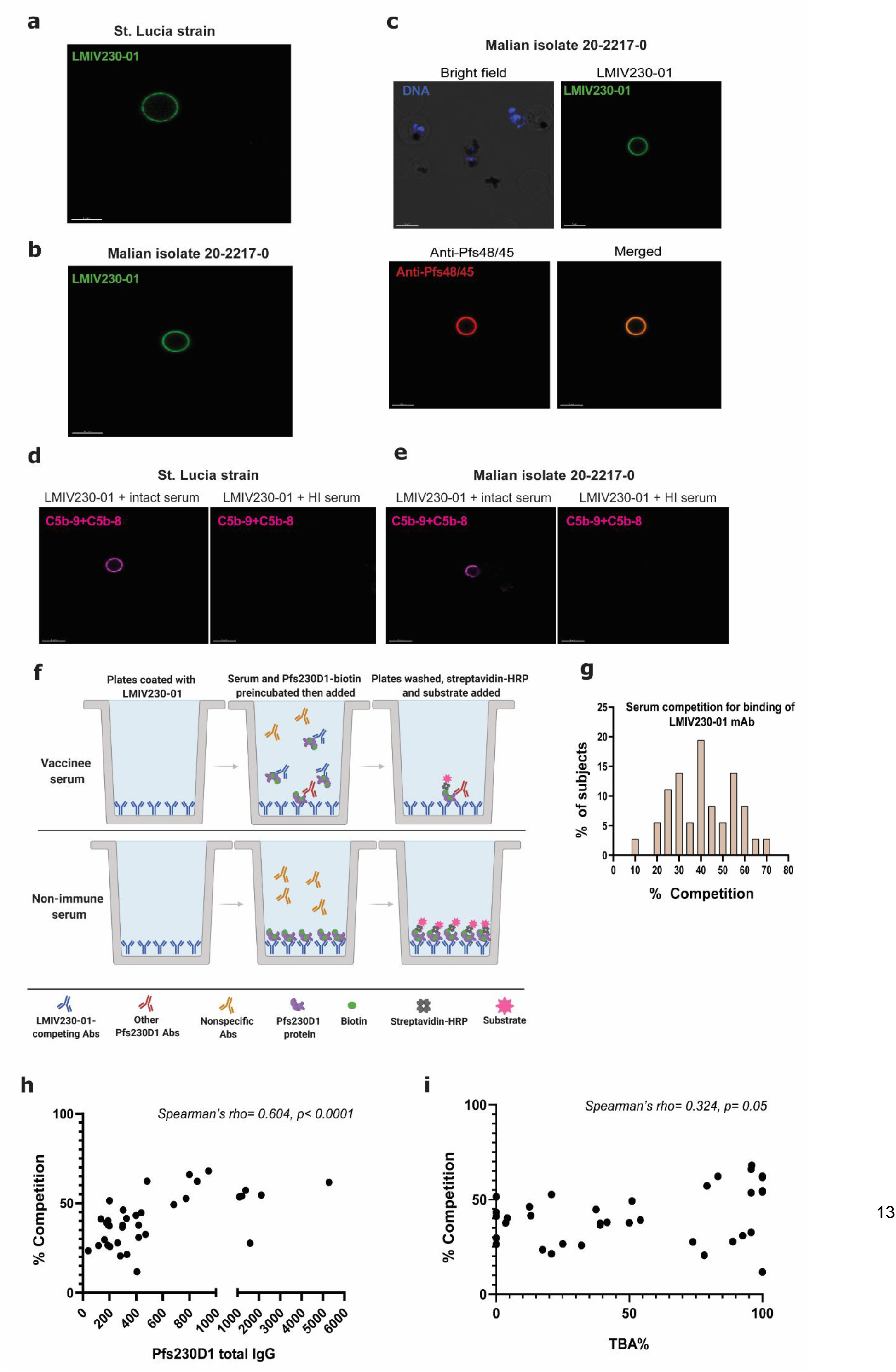
LMIV230-01 binds to heterologous *P. falciparum* strains and antisera from Pfs230D1 vaccinees vary widely in levels of antibody that compete with LMIV230-01 for binding. **a,** LMIV230-01 bound to gametes of *St. Lucia* parasite strain and **b,** of an isolate obtained from a Malian adult and adapted to culture. **c,** Murine anti-48/45 mAb confirms formation of gametes by Malian isolate and its signal colocalizes with LMIV230-01. “Merged” refers to combination of green and red channels. **d,** Membrane attack complex forms on gametes of St. Lucia strain and **e,** of a Malian isolate incubated with LMIV230-01 in the presence of intact but not heat-inactivated serum. Scale bars for all images in this panel: 5μM. **f,** Cartoon schematizing LMIV230-01 competition ELISA assay. **g,** Distribution of serum antibody levels that compete with LMIV230-01 for binding to Pfs230D1 in 36 subjects who received Pfs230D1-EPA vaccine. Values displayed represent mean from three independent experiments. **h,** Relationship of LMIV230-01-competing antibody levels to total Pfs230D1 antibody titers, or **i**, to serum functional activity (TBA, transmission blocking activity) measured by SMFA.

To assess the abundance of antibodies that share paratope specificity with LMIV230-01, we developed an ELISA assay to demonstrate the competition between post-vaccination sera (tested at a 1:2500 dilution) and LMIV230-01 for binding the vaccine antigen **(Figure 3f)**. Among subjects who received the vaccine, levels of competing antibody ranged from ~10-70% displacement of Pfs230D1 binding to LMIV230-01, with a normal distribution confirmed by Shapiro-Wilk test (p= 0.52) **(Figure 3g)**. Levels of competition strongly correlated with total Pfs230D1 IgG titers in sera (Spearman’s rho= 0.604, p<0.0001) **(Figure 3h**). Increasing levels of competing antibody also corresponded to serum functional activity measured by SMFA. Because serum TRA levels of vaccines were high with minimal variability ranging from 95-100% **(Extended Data Figure 14)**, our primary correlation analysis used TBA (Transmission Blocking Activity) which indicates the % reduction in the proportion of infected mosquitoes, a high bar for TBV activity generally seen only when TRA is very high. Correlation analyses showed that % serum competition was related to TBA (Spearman’s rho= 0.324, p= 0.05) **(Figure 3i)**, suggesting that antibodies that compete for the LMIV230-01 epitope play an important role in serum functional activity. This result, however, does not exclude the possible role of antibodies that do not compete with LMIV230-01 in mediating vaccine activity, and notably some sera with high TBA demonstrated low levels of LMIV230-01 competing antibodies.

Altogether, our data confirm that vaccination with TBV can elicit strong neutralizing antibodies, capable of binding to gametocytes, gametes, and zygotes, and of impairing fertilization in the mosquito. Due to its complex domains and repeating motifs with numerous disulfide bonds, expression of full length Pfs230 has been difficult^22,23^. Preclinical studies of Pfs230 fragments have shown that immunization with recombinant domain 1 of Pfs230 (Pfs230D1), but not other domains, induces strong functional TRA in SMFA^3,22,24^.

Our data support further development of TBV strategies to induce potent antibody responses against mosquito sexual stage parasites.

## ACKNOWLEDGMENTS

This work was funded by the Intramural Research Program of the National Institute of Allergy and Infectious Diseases, National Institutes of Health. Calvin Eigsti provided support for single cell sorting; Sundar Ganesan for live cell imaging; Ashley McCormack and Emily Higbee for Standard Membrane Feeding Assays; and J. Patrick Gorres for proofreading and editing this manuscript. We are thankful to Jillian Neal and Robert Morrison for determining the Pfs230D1 sequence for the Malian *P. falciparum* isolate. JT and JDG were supported by the Swiss National Science Foundation (Ambizione-SCORE fellowship to JT: PZ00P3_161147; PZ00P3_183777). We thank the staff members of GM/CA beamline at the Advanced Photon Source, Argonne National Laboratory for beamline support.

## AUTHOR CONTRIBUTIONS

C.H.C. and P.E.D. conceived the single B cell sorting of Pfs230D1-specific B cells, V gene repertoire analyses, antibody generation, conventional and competitive ELISAs, Western blot, microscopy-based binding assays and in vitro and in vivo functional characterization of mAbs. W.K.T. and N.H.T. conceived the epitope binning and biophysical studies. C.H.C, W.K.T., N.H.T and P.E.D conceived the analysis of polymorphisms. C.H.C., W.K.T., M.B., J.R., A.S., T.A.S., W.P., X.H., B.B., O.M., B. J, M.S. and N.D.S. performed the experiments. M.E., C.H.C., and J.D.G. performed bioinformatic analyses. N.J.M., K.R., V.N., R.H., R.S. and D.N. generated recombinant Pfs230D1. I.S., J.J.T., J.V.R., J.T., J.H., M.B.S, J.R., N.H.T. and P.E.D. supervised the experiments and interpreted the data. C.H.C., W.K.T., N.H.T. and P.E.D. wrote the manuscript, with input from all authors.

## COMPETING INTERESTS

M.B.S, W.P., X.H., B.B., and M.E. declare competing financial interests as all are employees of iRepertoire Inc., and J.H. is co-founder and CEO. J.D.G. is an employee of Alchemab Therapeutics Limited.

## CODE AVAILABILITY

Code is available on request from the corresponding author.

## EXTENDED DATA - FIGURES

**Extended Data Fig. 1.**
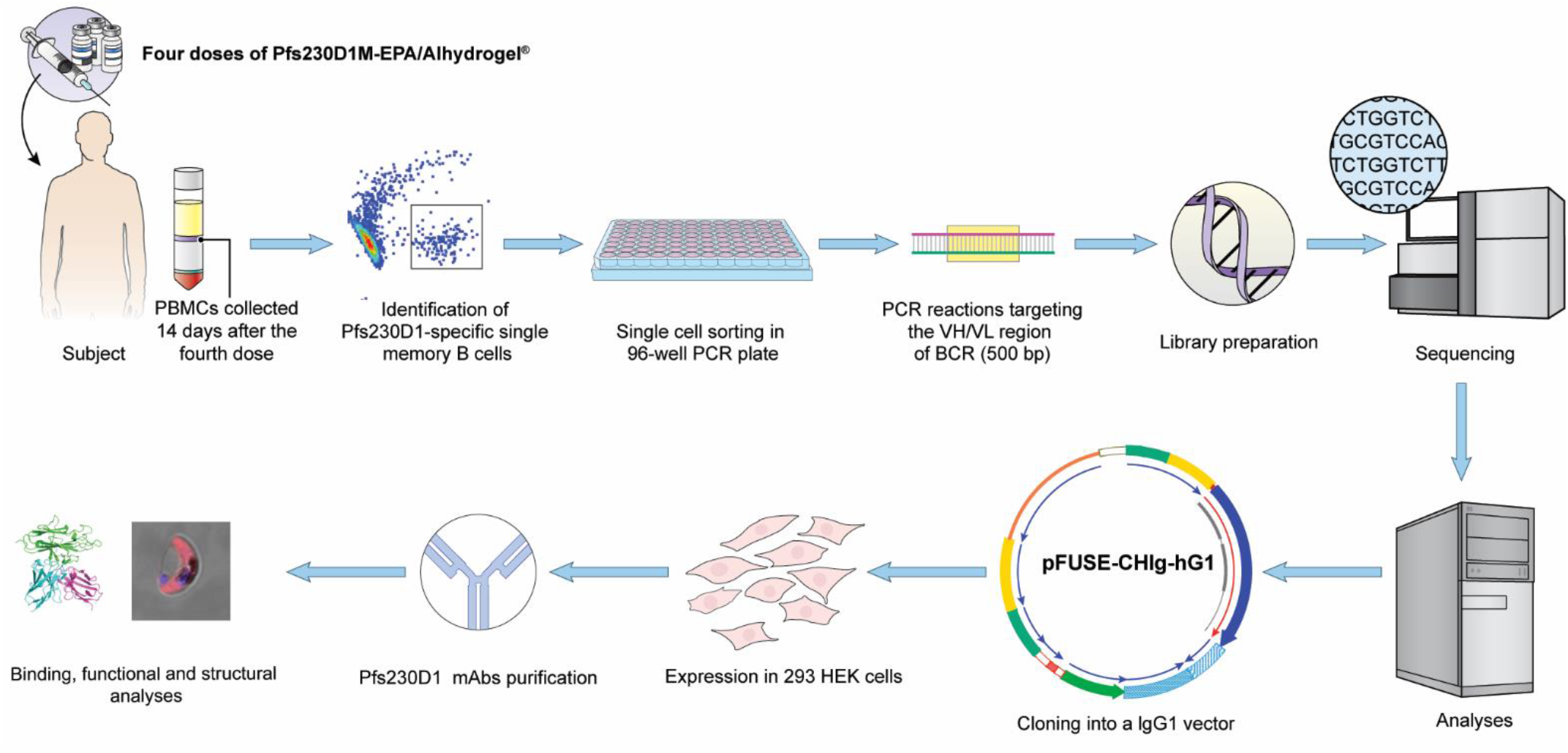
Experimental pipeline. Pfs230D1-specific single B cells were sorted from PBMCs of eight Malian adults who had been immunized with four doses of 40μg of Pfs230D1-EPA/Alhydrogel®. After extraction of single B cells, a 500 bp fragment of the BCR variable regions of VH/VL were amplified and sequenced. Matched VH/VL pairs that were identified in more than one B cell were preferentially selected for cloning in an IgG1 vector for expression in 293 HEK cells and subsequent analyses.

**Extended Data Fig. 2.**
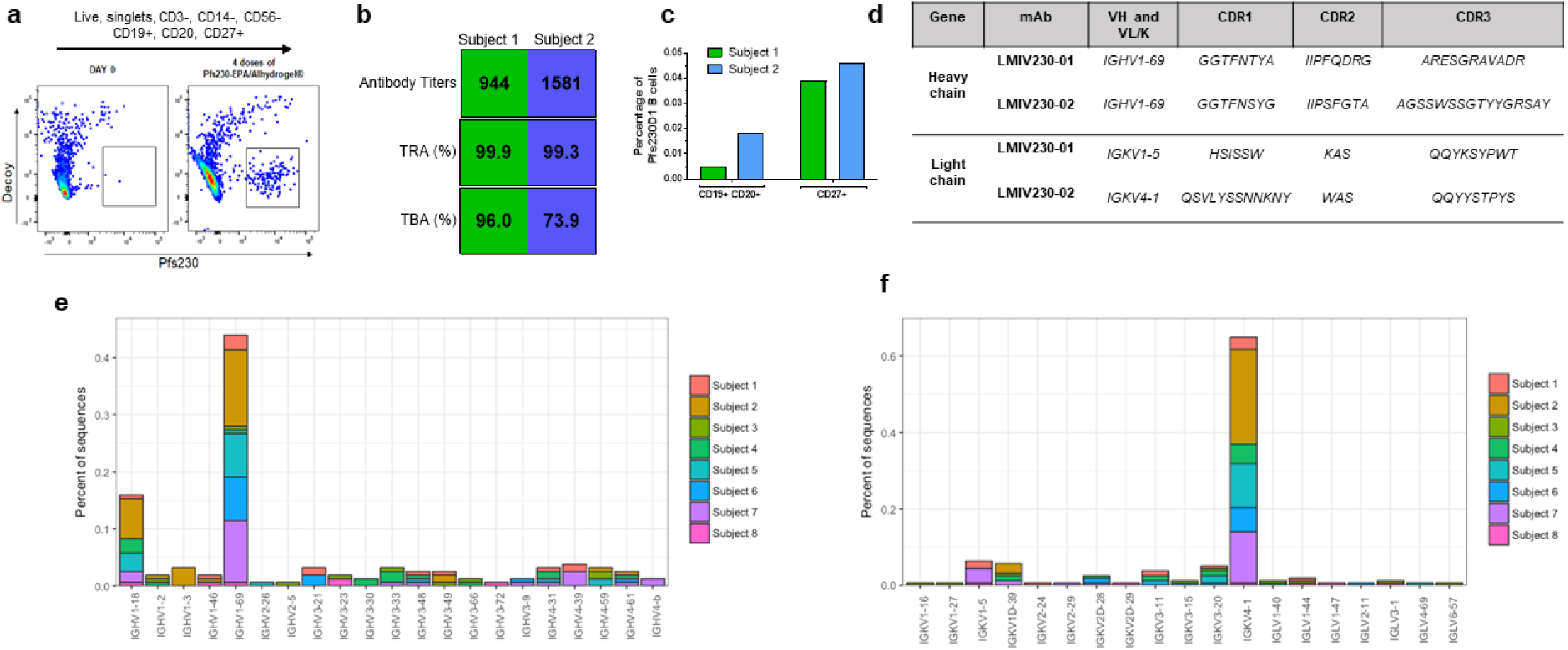
Pfs230D1-specific mAbs belong to the same heavy chain germline subgroup but differ for kappa chain. **a,** Sorted memory B cells were gated as live, single cells, excluded for CD3, CD14 and CD56, and gated on CD19+, CD20+, CD27+ cells. Then, a tetramer approach was used to select antigen-specific cells and reduce nonspecific binding. Cells binding to the decoy tetramer (BSA) were excluded and only those binding to Pfs230D1 were selected for sorting. **b,** Serum from each subject was used to measured antibody titers against Pfs230D1 and functional activity to reduce oocyst burden in Standard Membrane Feeding Assays (SMFA). TRA= Transmission Reducing Activity measured as the reduction in average oocyst count; TBA= Transmission Blocking Activity measured as the reduction in the proportion of infected mosquitoes. **c,** Proportion of memory B cells for each subject that are Pfs230D1-specific. **d,** Complementarity-determining regions (CDRs) of each sequence selected for mAb expression. **e,** IGKV4-1 germline (gene sequence in LMIV230-02) was the most frequent for the kappa chain genes. IGKV1-5 germline (gene sequence in LMIV230-01) was found in only three subjects **f,** Sequences related to germline 1-69 of the IGHV gene were the most frequently elicited in response to the vaccination.

**Extended Data Fig. 3.**
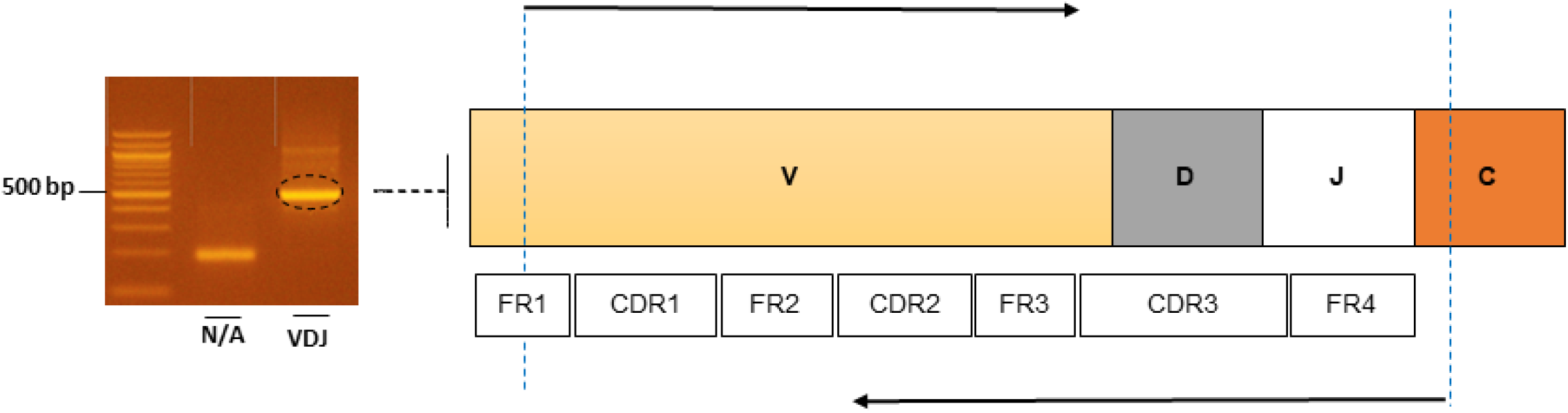
Amplification of V(D)J region. 500 bp fragment amplified from cDNA of sorted Pfs230D1-specific single B cell. This fragment was obtained using primers targeting the V(D)J region (iRepertoire Inc.).

**Extended Data Fig. 5.**
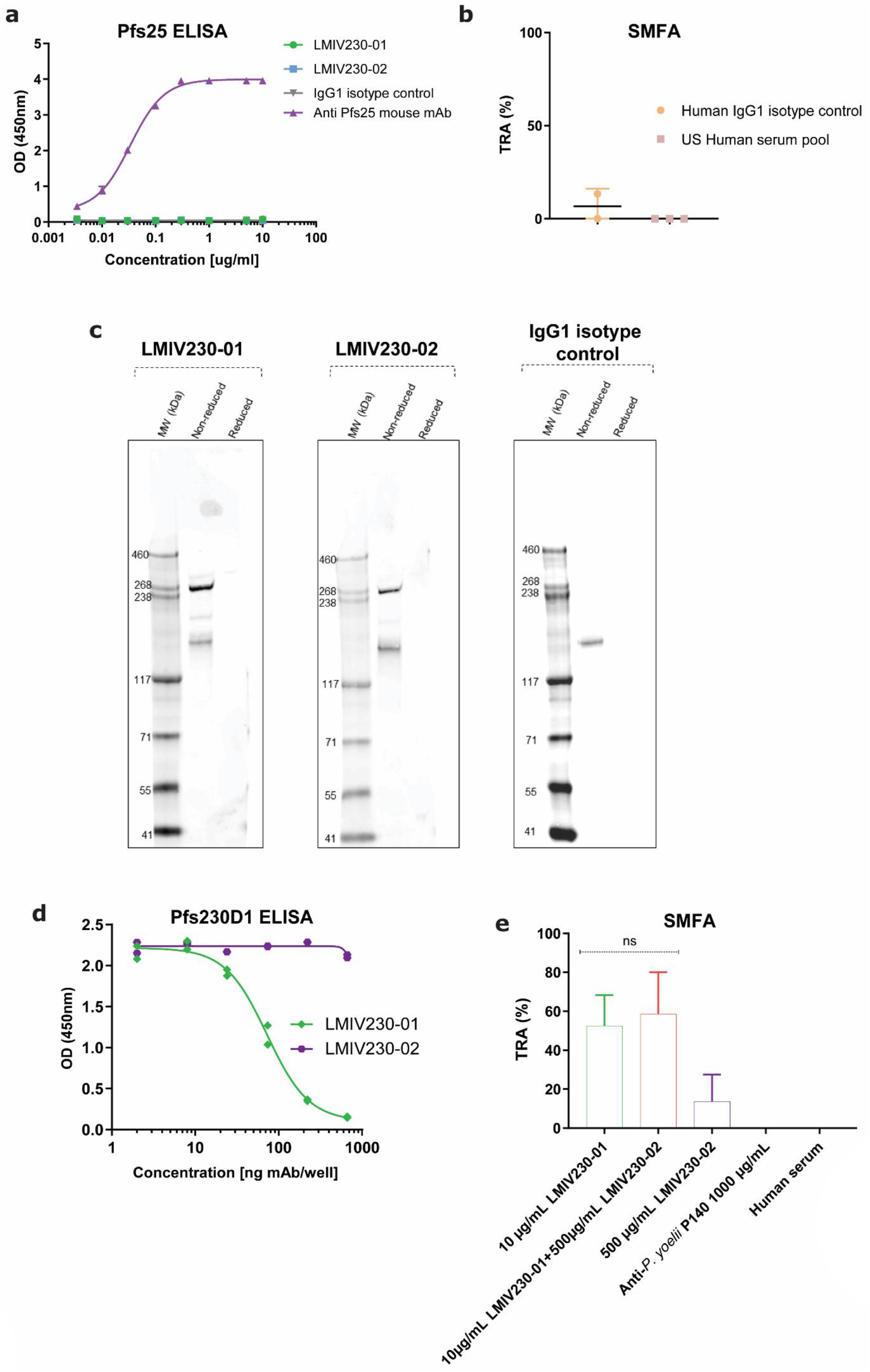
Additional binding and functional characterization of LMIV230-01 and −02. **a,** Both mAbs failed to bind to the ookinete protein Pfs25. **b,** Additional controls for the Standard Membrane Feeding Assay (SMFA). Human IgG1 isotype control was expressed using the same conditions as LMIV230-01 and −02 and was used in this assay at 1000μg/mL. Sixty microliters of undiluted human pooled serum obtained from US healthy donors were used as additional control. Values are shown as mean ± s.e.m. **c,** Full depiction of the Western blot gel displayed in Fig. 1g. **d,** The two mAbs do not compete for the same epitope in the recombinant Pfs230D1 protein, since unlabelled LMIV230-01 blocks binding of LMIV-230-01-HRP to immobilized Pfs230D1 but LMIV230-02 does not. **e,** Combination of LMIV230-01 and LMIV230-02 did not increase functional activity over LMIV230-01 alone. Control mosquitoes were fed with mouse IgG1 mAb targeting *P. yoelii* P140 protein, or with non-immune human serum.

[EXT. DATA FIGURE 6 WILL BE AVAILABLE IN THE PUBLISHED VERSION]

**Extended Data Fig. 7.**
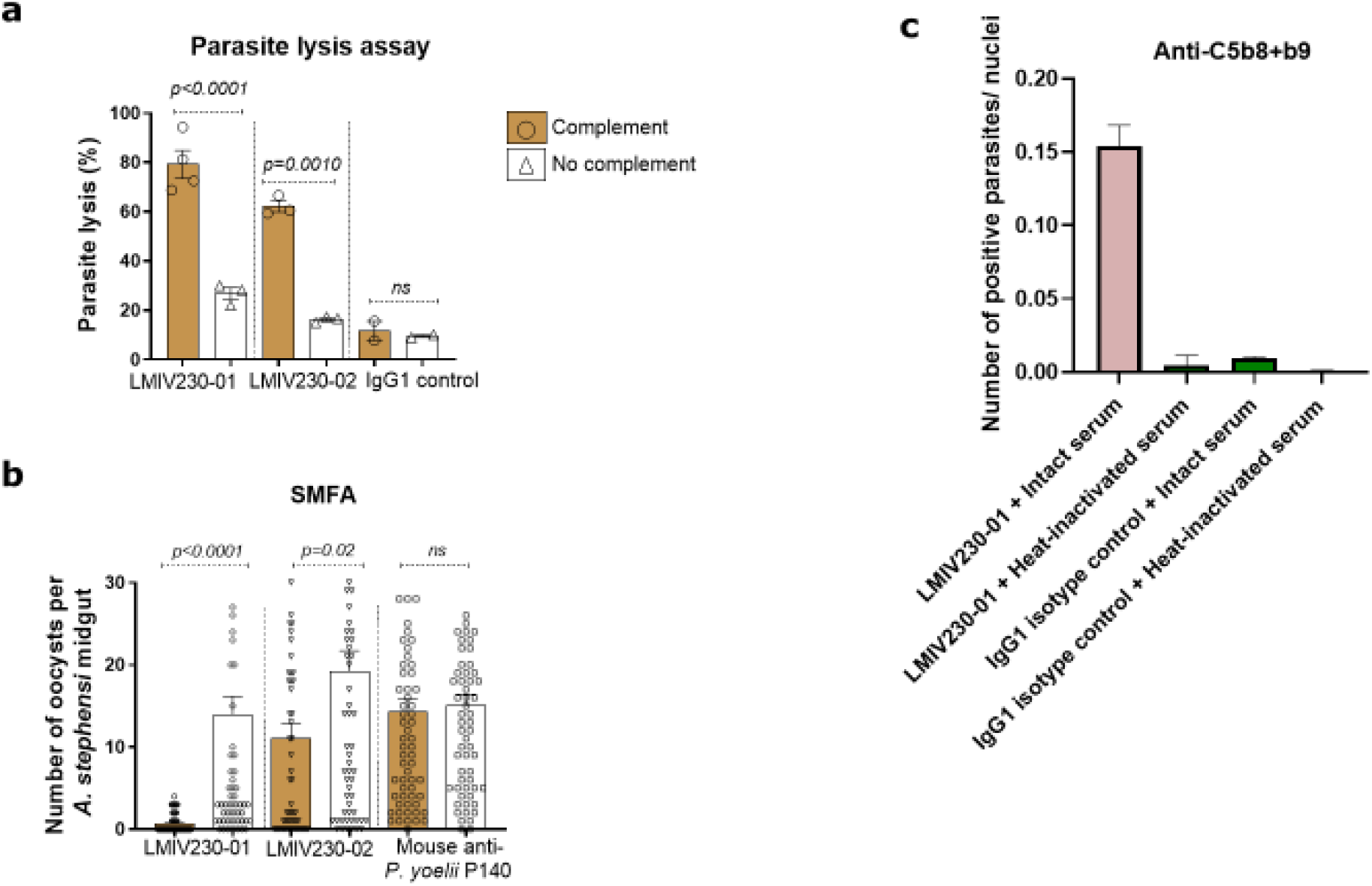
Pfs230 mAbs activity is complement-dependent and LMIV230-01 competing antibodies are acquired at varying levels by vaccinees. **a,** Activity of LMIV230-01 and LMIV230-02 is complement-dependent in the vitro lysis assay and **b,** in the vivo mosquito feeding assay. **c,** Membrane attack complexes (MAC) on parasites were detected using an Alexa 488-labeled antibody that recognizes the assembled MAC complex (anti C5b-9+ C5b-8). Gametes incubated with LMIV230-01 and intact serum produced MAC-positive parasites. Heat-inactivating serum to degrade the heat-labile components of the complement pathway eliminated deposition of MAC on gametes. MAC-positive *P. falciparum* strain NF54 gametes were enumerated in a large, tiled confocal image and normalized to the number of Hoechst-stained nuclei.

[EXT. DATA FIGURES 8-13 WILL BE AVAILABLE IN THE PUBLISHED VERSION

**Extended Data Fig. 14.**
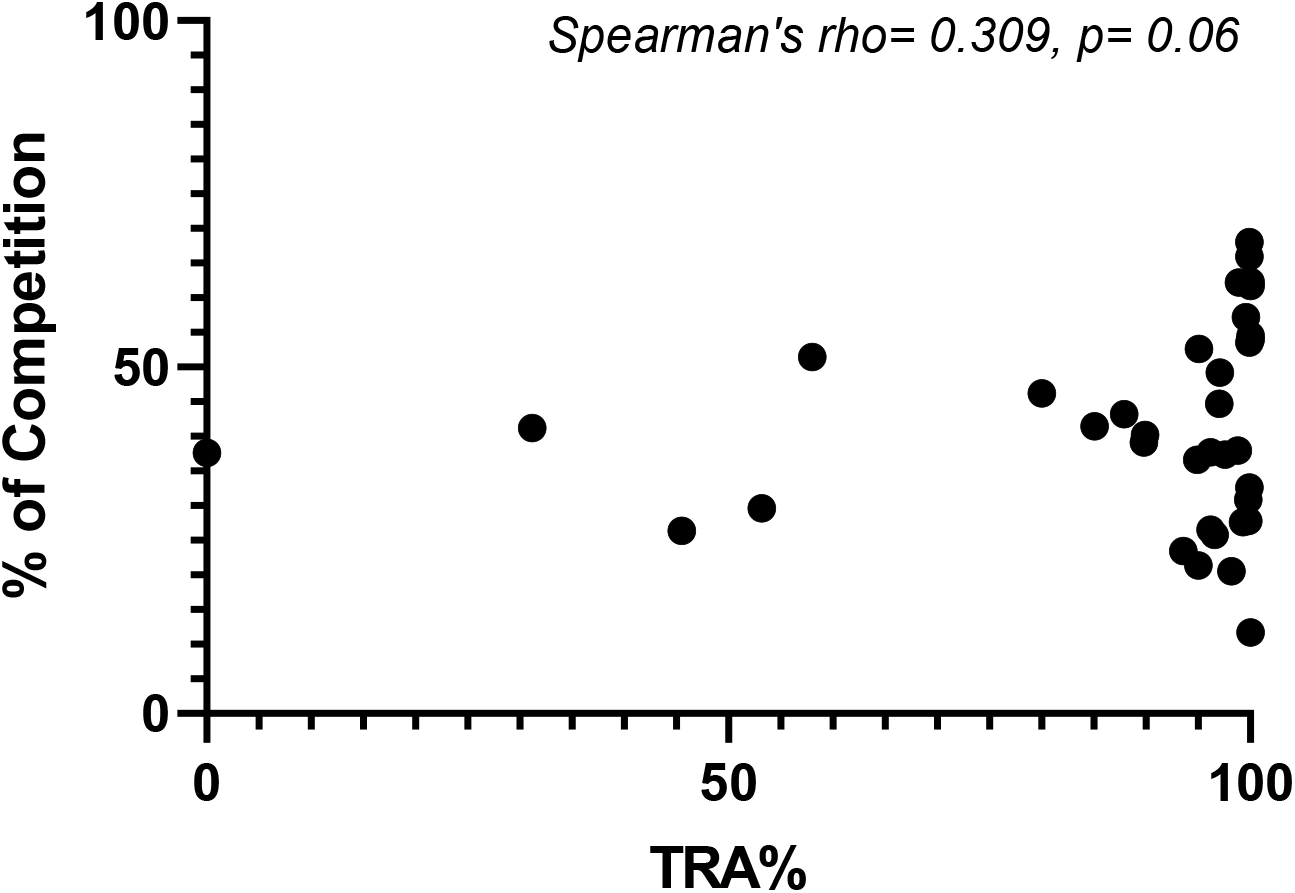
Correlation between levels of LMIV230-01 competing antibodies and Transmission-Reducing Activity (TRA) measured in SMFA.

## EXTENDED DATA – TABLES

**Extended Data Table 1.** Antibody titers and functional activity of sera from the eight subjects whose sequences were analyzed in this study. TRA= Transmission-reducing activity. TBA=Transmission blocking activity.

| Subject ID | Antibody titers | TRA (%) | TBA (%) |
| --- | --- | --- | --- |
| 1 | 944 | 99.9 | 96 |
| 2 | 1581 | 99.3 | 73.9 |
| 3 | 2115 | 100 | 100 |
| 4 | 1382 | 99.6 | 79.2 |
| 5 | 2100 | 100 | 100 |
| 6 | 5277 | 100 | 100 |
| 7 | 800 | 99.9 | 95.8 |
| 8 | 774 | 95.1 | 20.8 |

**Extended Data Table 2.** Binding of mAbs LMIV230-01 and LMIV230-02 to Pfs230D1 using Biolayer Interferometry. Binding data for each mAb was fitted using a 1:1 binding model. The averages for two biological replicates, composed of three technical replicates each, are shown for both mAbs.

| | $K_D$<br>($\times 10^{-10} \pm \text{SEM M}$) | $k_a$<br>($\times 10^5 \pm \text{SEM 1/Ms}$) | $k_{dis}$<br>($\times 10^{-4} \pm \text{SEM 1/s}$) | N |
| --- | --- | --- | --- | --- |
| **LMIV230-01** |  |  |  |  |
| **Biological Replicate 1** | $1.58 \pm 0.77$ | $1.71 \pm 0.06$ | $0.28 \pm 0.15$ | 3 |
| **Biological Replicate 2** | $2.06 \pm 0.99$ | $1.80 \pm 0.04$ | $0.37 \pm 0.18$ | 3 |
| **LMIV230-02** |  |  |  |  |
| **Biological Replicate 1** | $6.36 \pm 0.24$ | $7.67 \pm 0.21$ | $4.87 \pm 0.06$ | 3 |
| **Biological Replicate 2** | $4.27 \pm 0.22$ | $6.37 \pm 0.13$ | $2.71 \pm 0.10$ | 3 |

[EXT. DATA TABLES 3-7 WILL BE AVAILABLE IN THE PUBLISHED VERSION]

## Notes

### Summary of Updates

Malaria elimination requires tools that interrupt parasite transmission. Here, we characterized B cell receptor responses among Malian adults vaccinated against the first domain of the cysteine-rich 230kDa gamete surface protein Pfs2301-3 to neutralize sexual stage P. falciparum parasites and halt their further spread. We generated nine Pfs230 human monoclonal antibodies (mAbs). One mAb potently blocked transmission to mosquitoes in a complement-dependent manner and reacted strongly to gamete surface while eight mAbs showed only low or no blocking activity. This study provides a rational basis to improve malaria vaccines and develop therapeutic antibodies for malaria elimination.

